# NanoDam identifies novel temporal transcription factors conserved between the *Drosophila* central brain and visual system

**DOI:** 10.1101/2021.06.07.447332

**Authors:** Jocelyn L.Y. Tang, Anna E. Hakes, Robert Krautz, Takumi Suzuki, Esteban G. Contreras, Paul M. Fox, Andrea H. Brand

## Abstract

Temporal patterning of neural progenitors is an evolutionarily conserved strategy for generating neuronal diversity. Type II neural stem cells in the *Drosophila* central brain produce transit-amplifying intermediate neural progenitors (INPs) that exhibit temporal patterning. However, the known temporal factors cannot account for the neuronal diversity in the adult brain. To search for new temporal factors, we developed NanoDam, which enables rapid genome-wide profiling of endogenously-tagged proteins *in vivo* with a single genetic cross. Mapping the targets of known temporal transcription factors with NanoDam identified Homeobrain and Scarecrow (ARX and NKX2.1 orthologues) as novel temporal factors. We show that Homeobrain and Scarecrow define middle-aged and late INP temporal windows and play a role in cellular longevity. Strikingly, Homeobrain and Scarecrow have conserved functions as temporal factors in the developing visual system. NanoDam enables rapid cell type-specific genome-wide profiling with temporal resolution and can be easily adapted for use in higher organisms.

## Introduction

The nervous system is generated by a relatively small number of neural stem cells (NSCs) and progenitors that are patterned both spatially and temporally (Holguera and Desplan, 2018). Spatial patterning confers differences between populations of NSCs, while changes in gene expression over time direct the birth order and subtype identity of neuronal progeny. Temporal transcription factor cascades determine neuronal birth order in the *Drosophila* embryonic central nervous system (CNS) and the larval central brain and optic lobe (Doe, 2017). In the central brain, Type II NSCs generate transit-amplifying intermediate neural progenitors (INPs), which divide asymmetrically to self-renew and generate daughter cells (ganglion mother cells or GMCs) in a manner analogous to human outer radial glial cells (Bello et al., 2008; Boone and Doe, 2008; Bowman et al., 2008; Hansen et al., 2010). GMCs in turn undergo a terminal cell division, generating neurons or glial cells that contribute to the adult central complex (Bayraktar and Doe, 2013; Bayraktar et al., 2010; Izergina et al., 2009; Viktorin et al., 2011). The sequential divisions of INPs increase the quantity of neurons, which in turn creates a platform for generating wider neuronal diversity: eight Type II NSCs in each brain lobe give rise to at least 60 different neuronal subtypes. The tight control of progenitor temporal identity is crucial for the production of neuronal subtypes at the appropriate time and in the correct numbers.

The INPs produced by the six dorsal-medial Type II lineages (DM1-6) sequentially express the temporal transcription factors Dichaete (D, a member of the Sox family), Grainy head (Grh, a Grh/CP2 family transcription factor) and Eyeless (Ey, a homologue of Pax6) (Fig. 1A) (Bayraktar and Doe, 2013). These temporal factors were discovered initially by screening Type II lineages for restricted expression of neural transcription factors, using 60 different antisera (Bayraktar and Doe, 2013). This non-exhaustive approach was able to find a fraction of the theoretically necessary temporal factors, leaving the true extent of temporal regulation and the identity of missing temporal factors open. Furthermore, the cross-regulatory interactions predicted in a temporal cascade, in which each temporal transcription factor activates expression of the next temporal factor and represses expression of the temporal factor preceding it, are not fulfilled solely by D, Grh and Ey (Baumgardt et al., 2009; Brody and Odenwald, 2000; Cleary and Doe, 2006; Grosskortenhaus et al., 2005; 2006; Isshiki et al., 2001; Kambadur et al., 1998; Novotny et al., 2002; Pearson and Doe, 2003; Tran and Doe, 2008; Li et al., 2013).

**Figure 1:**
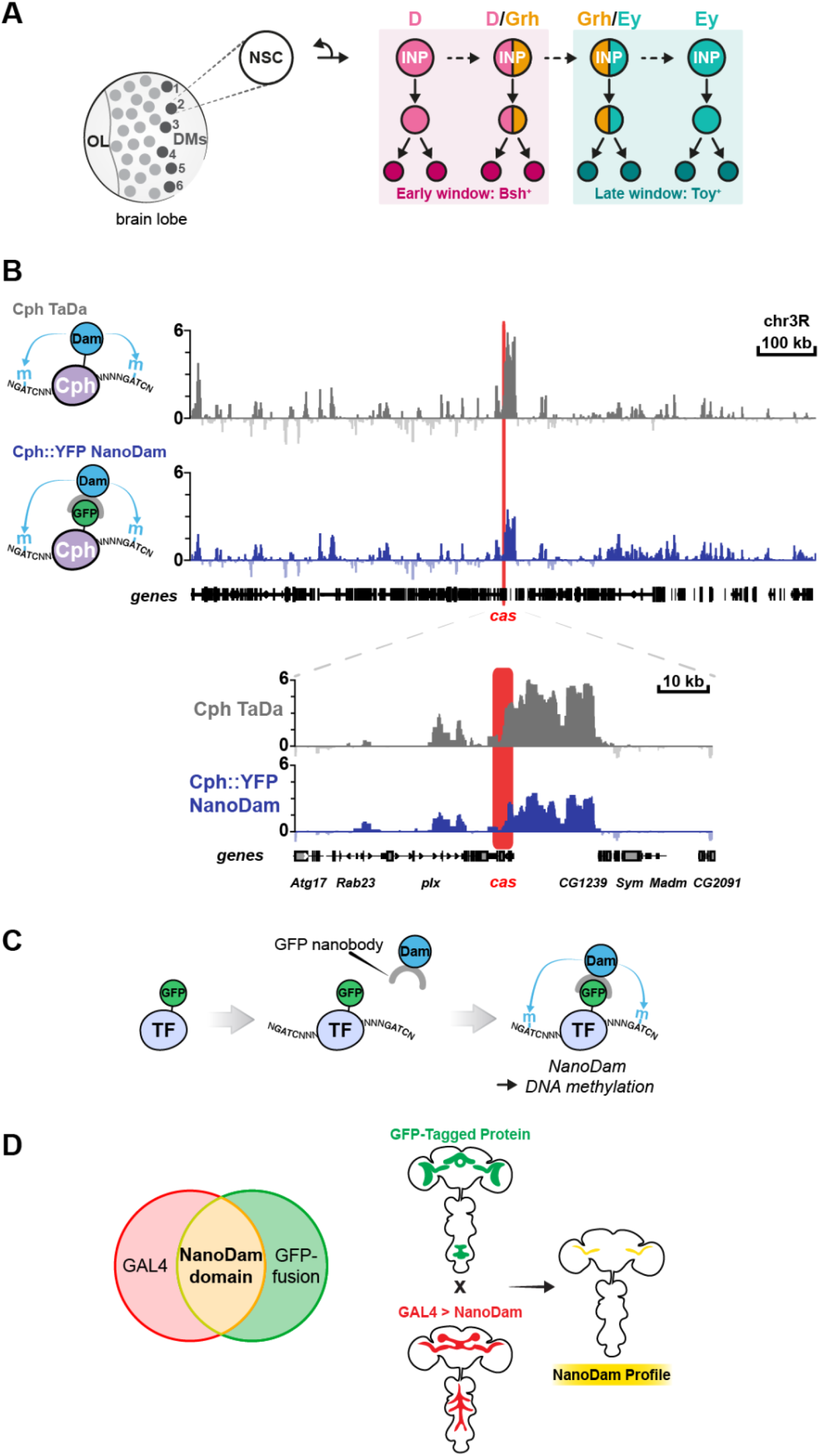
NanoDam profiles the genome-wide binding sites of GFP-tagged transcription factors in their endogenous temporal windows. (**A**) In the *Drosophila* larval CNS, the six dorsal-medial (DM) Type II NSCs in each brain lobe generate INPs that sequentially express D, Grh and Ey. The expression of D/Grh vs Grh/Ey defines early and late INP temporal windows and contributes to neuronal diversity. (**B**) Comparison of Cph TaDa and NanoDam (using Cph::YFP) binding across a region of chromosome 3R. Note that *cas* (highlighted in red), which we previously identified as a target of Cph using TaDa, is detected with both techniques. **(C**) To create NanoDam, a GFP nanobody was fused to the Dam protein. The nanobody can bind to the GFP-tagged transcription factor (TF) and so recruits Dam to the TF binding sites, resulting in genome-wide GATC methylation. (**D**) NanoDam allows for increased spatial resolution of profiling.

Three further factors contribute to INP temporal progression, but they are expressed broadly rather than in discrete temporal windows: Osa, a SWI/SNF chromatin remodelling complex subunit, and two further transcription factors, Odd-paired (Opa) (Abdusselamoglu et al., 2019) and Hamlet (Ham) (Eroglu et al., 2014). Therefore, there must exist other transcription factors that are expressed in defined temporal windows and exhibit the regulatory interactions expected in a temporal cascade. We postulated that other temporal factors, that contribute to generating the diversity of neuronal subtypes arising within each INP lineage, remain to be identified.

Given the feed-forward and feed-back transcriptional regulation previously observed in temporal transcription cascades, we surmised that novel temporal factors would be amongst the transcriptional targets of D, Grh or Ey. Therefore, we devised a novel approach, NanoDam, to identify the genome-wide targets of transcription factors within their normal expression windows *in vivo* without cell isolation, cross linking or immunoprecipitation. Temporal factors are expressed transiently in a small pool of rapidly dividing progenitor cells. NanoDam provides a simple, streamlined approach to obtain genome-wide binding profiles in a cell-type-specific and temporally restricted manner.

Using NanoDam, we identified the transcriptional targets of D, Grh and Ey in INPs and, by performing single cell RNA sequencing, determined which of the directly bound loci were activated or repressed. Next, we assessed which of the target loci encoded transcription factors and whether these were expressed in restricted temporal windows within INPs. We surveyed where in the INP transcriptional cascade these factors acted and ascertained whether they cross regulate the expression of other temporal transcription factor genes, as expected for temporal factors. Finally, we showed that the newly discovered temporal factors play the same roles, and exhibit the same cross regulatory interactions, in the temporal cascade in the developing visual system. This is particularly striking as the INPs and the NSCs of the developing optic lobe have different cells of origin and yet the mechanism they use to generate neural diversity is conserved.

## Results

### NanoDam for genome-wide binding profiling of endogenously tagged chromatin-binding proteins

To identify novel temporal transcription factors, we set out to profile the genome-wide binding targets of D, Grh and Ey, within their normal temporal windows. To this end we devised a novel approach, NanoDam, to profile binding of endogenously tagged proteins *in vivo* without cell isolation, cross linking or immunoprecipitation. NanoDam capitalizes on our Targeted DamID technique (TaDa), in which the DNA- or chromatin-binding protein of interest is fused to an *E. coli* Dam methylase (Marshall and Brand, 2015; Marshall et al., 2016; Southall et al., 2013). As in DamID, when the Dam fusion protein interacts with the genome, it methylates adenine within the sequence GATC (van Steensel and Henikoff, 2000; van Steensel et al., 2001). Endogenous adenine methylation is extremely rare in eukaryotes (Koziol et al., 2016; Wu et al., 2016; Zhang et al., 2015), such that the genomic targets of the Dam-fusion protein can be identified readily by mapping adenine methylation in the genome (see for example, (Marshall et al., 2016)). Targeted DamID enables cell type-specific genome-wide profiling *in vivo*, using the GAL4 system (Brand and Perrimon, 1993), while avoiding the potential toxicity resulting from expression of high levels of the Dam methylase (Southall et al., 2013). For each TaDa experiment, however, a transgene must first be generated encoding the Dam methylase fused to a candidate protein. The transgene is then ectopically expressed, albeit at very low levels, driven by a cell type specific GAL4 driver (Southall et al., 2013).

In designing NanoDam, we sought to benefit from all of the advantages of TaDa whilst, first, bypassing the need to generate recombinant transgenes and, second, assessing genome-wide binding only when and where the DNA- or chromatin-binding protein is normally expressed. NanoDam recruits the Dam methylase to endogenously-tagged proteins, using nanobodies as targeting agents. Nanobodies are recombinant antibody fragments derived from the variable region of heavy chain antibodies present in camelid species (Muyldermans, 2001). We fused the Dam methylase to a nanobody recognizing GFP in order to direct the Dam methylase-nanobody fusion to any GFP-tagged protein and enable genome wide profiling of the tagged protein (Fig. 1C). Different nanobodies can be used to target proteins with specific post-translational modifications, or with tags other than GFP (Brand lab, unpublished).

We fused nanobody, vhhGFP4, which recognizes GFP and a number of its variants, including eGFP, YFP, CFP, BFP and Venus (Caussinus et al., 2012), to the C-terminus of Dam methylase, adding an intervening nuclear localisation sequence (Fig. S1A), and used TaDa (Southall et al., 2013) to drive low levels of tissue specific expression. With TaDa, a bicistronic message is transcribed: a primary open reading frame (ORF1; here mCherry) followed by two TAA stop codons and a single nucleotide frameshift upstream of a secondary open reading frame, in this case the coding sequence of the Dam-nanobody fusion protein (ORF2). Translation of this bicistronic message results in expression of ORF1, as well as extremely low levels of expression of the Dam-nanobody fusion protein (ORF2) due to rare ribosomal re-entry and translational re-initiation.

To assess the efficacy of NanoDam, we profiled binding of the *Drosophila* homologue of CTIP^1/2^/Bcl11^a/b^, which we call Chronophage (Cph). The *cph* locus had been tagged by insertion of a YFP protein trap (Cph::YFP) (Lowe et al., 2014), resulting in Cph-YFP expression from its own promoter in the embryonic central nervous system (PMF and AHB, unpublished). We drove expression of NanoDam in embryonic NSCs with *worniu*-GAL4 and compared our results to those we obtained using TaDa. NanoDam for Cph::YFP and TaDa for Cph, were performed under the same conditions (see Methods). NanoDam accurately reproduced the binding profiles obtained with TaDa, as exemplified at the *castor* locus (Fig. 1B), demonstrating the reliability of the approach.

NanoDam enables any endogenously-tagged factor to be profiled after a single genetic cross without cell isolation, cross linking or immunoprecipitation. Importantly, NanoDam profiles genome-wide binding of proteins only within the cells where they are normally expressed. By restricting expression of the NanoDam construct using different GAL4 drivers, the genome wide binding pattern of any tagged factor can be assessed in a defined subset of its endogenous expression pattern. Genomic DNA methylation occurs only in cells that express both the NanoDam construct (under the control of GAL4) and the endogenously-tagged protein (under the control of its own regulatory elements) (Fig. 1D). This is particularly important when profiling proteins that are expressed in only a subset of a cells within a lineage, a fact we sought to exploit for identifying temporal transcription factors. Therefore, we were confident that NanoDam could be used to profile the genome-wide occupancy of the temporal transcription factors D, Grh and Ey.

### NanoDam reveals combinatorial binding patterns of the INP temporal factors

We expressed NanoDam in the INPs using *D*-GAL4 (GMR12E09-GAL4 (Bayraktar and Doe, 2013). We crossed *D-GAL4*; *UAS-NanoDam* to flies expressing endogenously tagged D-GFP, Grh-GFP or Ey-GFP (Kudron et al., 2018). NanoDam profiles binding only in a subset of the *D*-GAL4 expression pattern, in cells that also express the endogenously tagged transcription factor (Fig. 2A). We expressed NanoDam for 14 hours at the end of the third larval instar stage. Genomic DNA was extracted from approximately 50 dissected brains per sample and processed as described previously for TaDa in order to generate libraries of fragments corresponding to transcription factor bound regions (Marshall et al., 2016; Southall et al., 2013).

**Figure 2:**
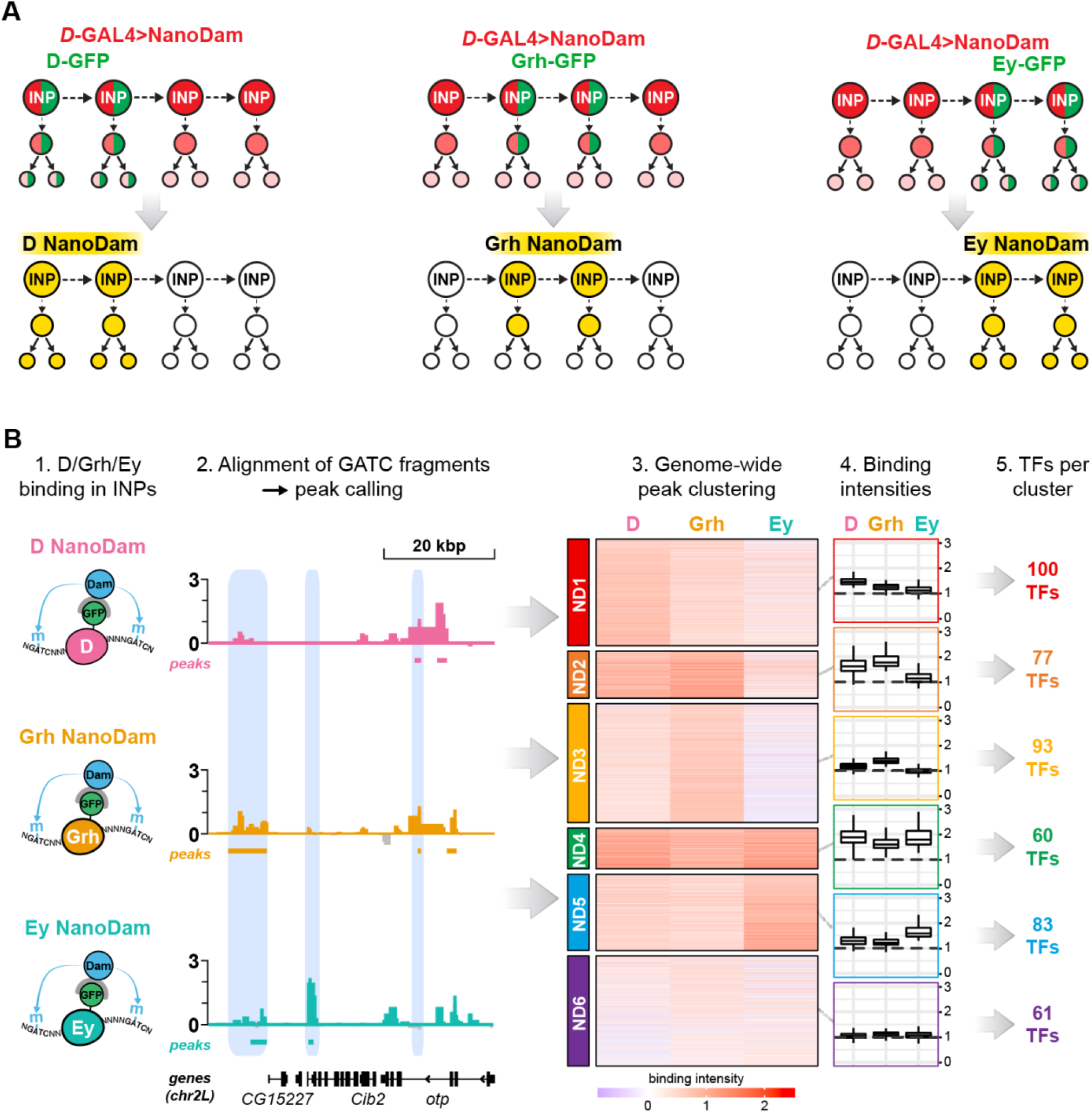
NanoDam for INP temporal factors D, Grh and Ey. (**A**) Schematic showing the experimental setup for D, Grh and Ey NanoDam in INPs. *D*-GAL4 drives UAS-*NanoDam* (red) in all INPs of DM1-6 during late third instar larval stage. D-GFP, Grh-GFP and Ey-GFP (green) are expressed in a subset of INPs and progeny, restricting NanoDam binding to their respective temporal windows (yellow). Note that *D*-GAL4 expression extends beyond the endogenous temporal window of D. (**B**) NanoDam-derived binding intensities for D, Grh and Ey were aggregated for highly significant peaks identified by comparison with the *w*^*1118*^ control. Unsupervised clustering of the peaks according to these intensities identified six distinct combinations of D/Grh/Ey binding, denoted as ND1-6. Binding intensities are shown as z-scores for individual peaks in the heat maps. Average binding intensities across all peaks per NanoDam cluster are represented as box plots. To assign peaks to genes, peaks belonging to each cluster were assigned to the nearest transcriptional start site and the resulting gene lists were subsetted for TFs.

As a first step in searching for novel temporal transcription factors regulated by D, Grh or Ey, we compared the NanoDam peaks for D, Grh and Ey to each other throughout the genome and clustered them according to their aggregated binding intensities. Clustering revealed 6 different combinations of D, Grh and Ey binding in INPs (Fig. 2B and S1B). Peaks in cluster ND1 corresponded to strong D binding but minimal binding of Grh or Ey; ND2 peaks showed strong D and Grh binding; ND3, strong Grh binding; ND4, strong binding of D, Grh and Ey; ND5, Ey binding; ND6, minimal binding of all three transcription factors. This suggested complex regulatory relationships between these temporal factors and their target genes.

To determine the functional relevance of these ND clusters, we assigned the peaks within each cluster to the nearest transcriptional start sites of protein-coding genes. Within the lists we identified genes encoding transcription factors (Fig. 2B and S1C) and hypothesised that some of these might be INP temporal transcription factors whose expression is regulated by D, Grh or Ey.

### scRNA-seq of INPs and their progeny

To assess whether the D-, Grh- and Ey-bound genes are regulated by these temporal factors, we carried out single cell RNA sequencing (scRNA-seq) of INPs and their progeny. We drove expression of membrane-targeted RFP (Mellert and Truman, 2012) in INPs with *D*-GAL4 and dissected brains from wandering third instar larvae. The dissected brains were dissociated enzymatically and RFP-positive cells were isolated by FACS. We recovered 4,086 single cells from approximately 230 brains across two biological replicates (Fig. S2A) with 2,614 median genes detected per cell. We clustered the cells with Seurat (Butler et al., 2018; Stuart et al., 2019), generating 12 clusters that were visualised using a t-distributed stochastic neighbour embedding (t-SNE) plot (Fig. 3A) (Maaten and Hinton, 2008). Cluster identities were assigned based on known cell-type-specific markers: cluster 1 was designated as INPs (*dpn, wor*) (Fig. S2B); clusters 2-10 as INP progeny; clusters 2-4 as immature neurons (*nSyb* and high *prospero*), and 9-10 corresponding as glial cells (*repo*) (Fig. S2C).

**Figure 3:**
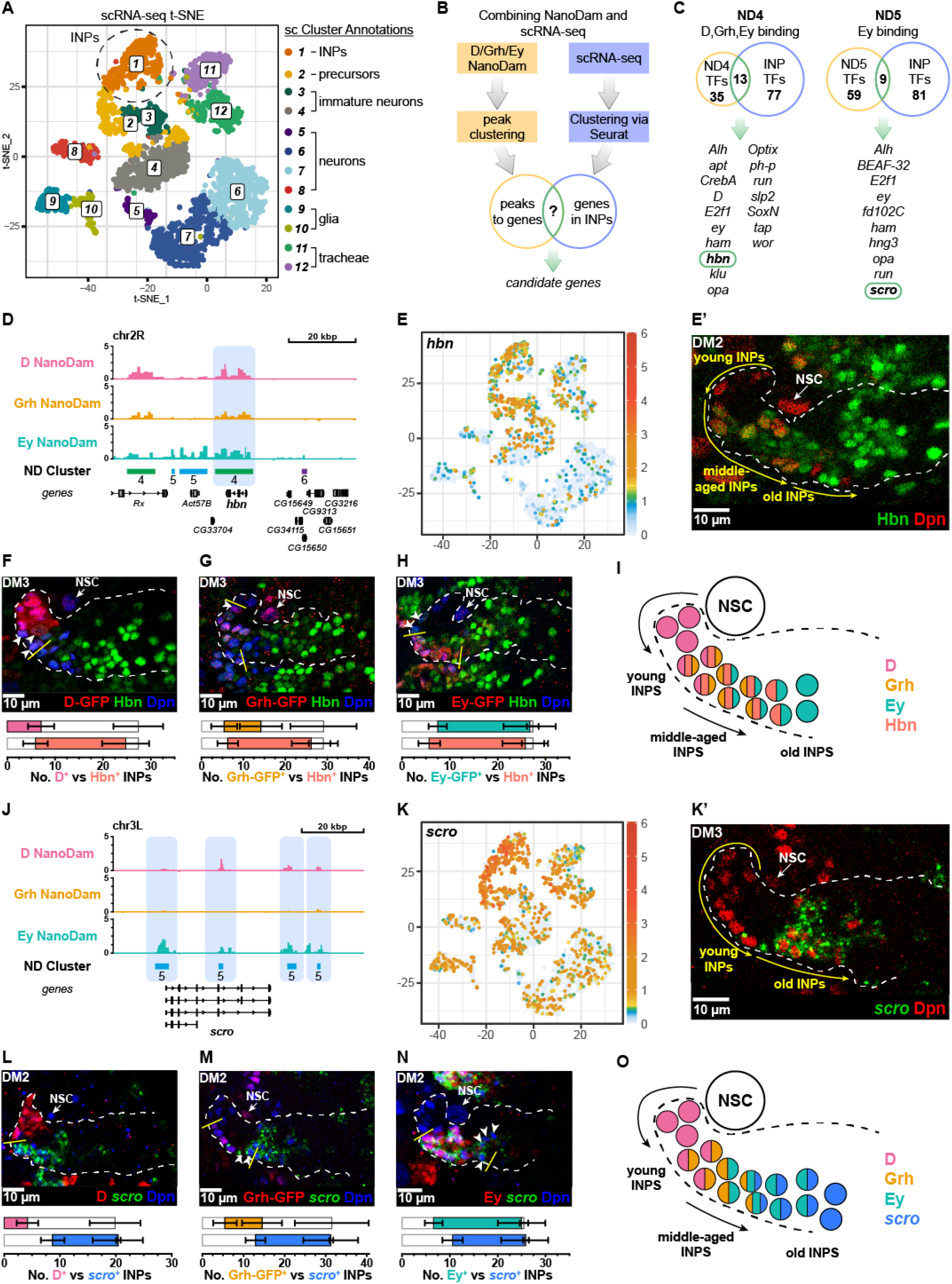
Combining NanoDam with scRNA-seq identifies *homeobrain*(*hbn*) and *scarecrow* (*scro*) as candidate temporal factors in INPs. (**A**) t-distributed stochastic neighbour embedding (t-SNE) visualisation of 4,086 sorted single cells coloured by cluster assignment and annotated based on previously known markers (see Fig. S2). (**B**) Schematic representation of the workflow used to identify candidate temporal factors. (**C**) Candidate temporal transcription factors *homeobrain* (*hbn*) and *scarecrow* (*scro*) (outlined in green) were significantly differentially expressed in INPs (based on the scRNA-seq data) and identified in NanoDam cluster 4 (ND4) or NanoDam cluster 5 (ND5). (**D**) NanoDam binding of D, Grh and Ey across the *hbn* gene locus and nearby regions. The ND6 peak across *hbn* is highlighted. **(E-E’**) *hbn* is expressed in INPs. (**E**) t-SNE plot coloured by *hbn* expression. (**E’**) Staining with an antibody raised against Hbn showed that Hbn (green) is not expressed in the Type II NSC (white arrow) but is expressed in INPs (Dpn^+^ (red)) and progeny (Dpn^-^). Hbn expression begins across a broad domain of middle-aged INPs but is absent from the oldest INPs. (**F**) Overlap of D-GFP (red) and Hbn (green) in INPs (Dpn^+^ (blue)). Hbn expression begins at the end of the D temporal window. Only the oldest D^+^ INPs express Hbn (arrowheads). Very few INPs expressed both D and Hbn and the few D^+^ Hbn^+^ INPs were found at the end of the D temporal window (Fig. S3A). (**G**) Hbn (green) overlaps with Grh^+^ (red) INPs. Arrowhead indicates a young Grh^+^ INP that does not express Hbn. (**H**) Hbn (green) expression begins before the Ey (red) window. Arrowheads indicate the youngest Hbn^+^ INPs that do not express Ey. (**I**) Summary schematic of the expression pattern of Hbn in relation to D, Grh and Ey. (**J**) NanoDam binding of D, Grh and Ey across the *scro* gene locus. The four ND5 peaks are highlighted. (**K-K’**) *scro* is expressed in INPs. (**K**) t-SNE plots coloured by *scro* expression. (**K’**) Fluorescence *in situ* hybridisation shows that *scro* (green) is not expressed in the Type II NSC (white arrow) or young INPs (Dpn^+^ (red)) but is expressed in old INPs (Dpn^+^). (**L**) *scro* (green) does not overlap with D (red) in INPs (Dpn^+^ (blue)). (**M**) The oldest Grh^+^ (red) INPs express *scro* (green) (arrowheads). (**N**) *scro* (green) expression begins after the activation of Ey (red) in INPs. The oldest *scro*^+^ INPs do not express Ey (arrowheads). (**O**) Summary schematic of the expression pattern of *scro* in relation to D, Grh and Ey. White dotted lines indicate *D*-GAL4>*mCD8-RFP* expression (which is expressed in INPs and their daughter cells) in single section confocal images. Brains were dissected at wandering third instar stage. *n* = 6 brain lobes.

As expected, all three temporal factors *D, grh* and *ey* were expressed in the INP cluster (Fig. S2D). We also observed significant expression of *grh* in clusters 11 and 12 (Fig. S2D) which, interestingly, did not show high levels of expression of brain-specific markers (Figs. S2B-C) but instead were enriched for tracheal gene expression (Fig. S2E). The expression of *grh* in these tracheal cell clusters is consistent with a previous study identifying a functional role for Grh in tracheal development (Yao et al., 2017). As such, we excluded clusters 11 and 12 from our analysis to focus solely on INPs and their progeny.

### Identifying new INP temporal factors

We determined which transcription factors in each NanoDam cluster were expressed in INPs by comparison with our scRNA-seq dataset (Fig. 3B and Table S1). In addition to transcription factors known to be involved in INP cell identity (*ase, dpn, wor*) and those shown previously to regulate temporal identity (*D, grh, ey, opa, ham*), we found several novel candidate temporal transcription factors. We focussed our attention on two factors that were expressed in a subset of INPs: the paired-like homeobox transcription factor *homeobrain* (*hbn*) (Walldorf et al., 2000) and the NK-2 homeobox transcription factor *scarecrow* (*scro*) (Zaffran et al., 2000) (Fig. 3C). *homeobrain* clusters in ND4 (strong binding of D, Grh and Ey; Fig. 3D), suggesting that multiple members of the temporal cascade may regulate its expression. *hbn* was also highly enriched in INPs according to our scRNA seq data (Fig. 3E). Examining Hbn expression in Type II lineages *in vivo* revealed that Hbn was expressed in middle-aged and old INPs but absent from NSCs and the youngest INPs (Figs. 3E’-I and Figs. S3A-D). In most Type II lineages, Hbn was expressed in INPs concomitantly with Grh (Fig. 3G and S3C) and prior to initiation of Ey expression (Figs. 3H and S3D). Hbn expression was maintained throughout most of the Ey temporal window, except in the oldest INPs in the DM2 and DM3 lineages (Fig. S3D). Therefore, Hbn expression defines a new temporal window extending from the end of D expression through the Ey expression window. *scro* clusters in ND5 (bound strongly by Ey and weakly by D, but not bound by Grh) (Fig. 3J) and is highly enriched in INPs (Fig. 3K). We assayed *scro* expression in Type II lineages *in vivo* by fluorescent *in situ* hybridization chain reaction (HCR) (Choi et al., 2014; 2016; 2010; 2018). *scro* was expressed in the oldest INPs in all DM lineages and absent from Type II NSCs and young INPs (Fig. 3K’). *scro* was never co-expressed with D (Fig. 3L and S3E) and only the oldest Grh^+^ INPs expressed *scro* (Fig. 3M and S3F). In all lineages (DM1-6) *scro* expression began after Ey and was maintained into the oldest INPs (Fig. 3N and S3G). Therefore, *scro* expression defines the latest temporal window (Fig. 3O).

### The middle-aged factor Hbn displays classic temporal factor interactions with Ey and *scro*

Next, we investigated the regulatory interactions between the known INP temporal factors and Hbn and Scro. Hbn is expressed in middle-aged INPs (Fig. 3I). Ey binds the *hbn* locus and is expressed in INPs after Hbn. Given that temporal factors are thought to repress transcription of the factor that precedes them, we tested whether Ey repressed Hbn expression. We found that ectopic expression of Ey in all INPs resulted in a significant reduction in Hbn expression (Fig. 4A), whereas knocking down *ey* expression in INPs (Fig. S4A) extended the Hbn window into the oldest INPs without affecting early Hbn expression (Fig. 4B and S4B). This was true of all lineages except DM5, in which the oldest INPs remained Hbn^-^ (Figs. S4B-B’). Knocking down *ey* with RNAi also increased the number of INPs in all DM lineages, as has been reported previously (Bayraktar and Doe, 2013). Therefore, Ey terminates Hbn expression in old INPs (Fig. 4H), a regulatory interaction characteristic of temporal factors.

**Figure 4:**
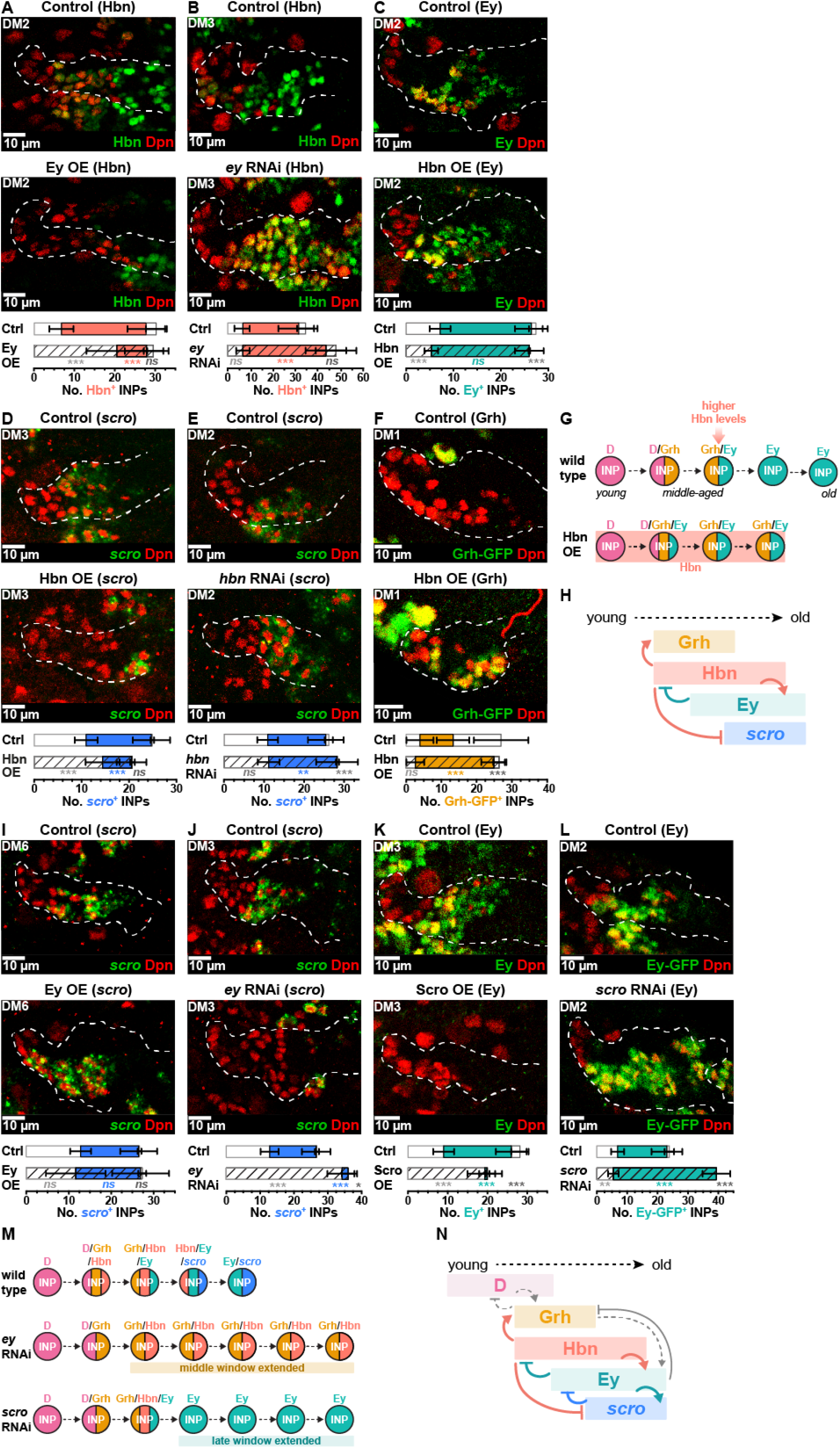
Homeobrain and scarecrow exhibit regulatory relationships typical of temporal factors. (**A**) Ey overexpression represses Hbn (green) in INPs (Dpn^+^ (red)). Mann-Whitney Test p<0.001, ***; p<0.001, ***; p=0.08, ns. (**B**) *ey* RNAi results in more Hbn^+^ (green) in INPs (Dpn^+^ (red)). Control is *w*^*1118*^. Mann-Whitney Test p=0.93, ns; p<0.001, ***; p=0.08, ns. (**C**) Hbn overexpression precociously activates Ey (green) in INPs (Dpn^+^). Mann-Whitney Test p<0.001, ***; p=0.15, ns; p<0.001, ***. (**D**) Hbn overexpression represses *scro* (green). Mann-Whitney Test p<0.001, ***; p<0.001, ***; p=0.73, ns. (**E**) Hbn knockdown leads to an increase in *scro* (green). Mann-Whitney Test p=0.83, ns; p=0.002, **; p=0.0005 ***. (**F**) Hbn overexpression leads to ectopic activation of Grh (green) in INPs (Dpn^+^ (red)). Control image is a projection over 8 µm and Hbn OE is a projection over 29 µm. Mann-Whitney Test p=0.13, ns; p<0.001, ***; p<0.001, ***. (**G**) Schematic summary of the phenotype of Hbn overexpression. Note in wildtype the strength of Hbn expression is higher later in the cascade. (**H**) Summary of the regulatory relationships of Hbn in the temporal cascade. (**I**) Ey OE precociously activates *scro* (green) in INPs (Dpn^+^ (red)). Mann-Whitney Test p=0.12, ns; p=0.44, ns; p=0.60. See Fig. S4G. (**J**) *ey* RNAi leads to loss of *scro* (green) in INPs (Dpn^+^ (red)). Mann-Whitney Test p<0.001, ***; p<0.001, ***; p=0.02, *. (**K**) Overexpression of Scro leads to loss of Ey (green) in INPs (Dpn^+^ (red)). Mann-Whitney Test p<0.001, *** for all. (**L**) *scro* RNAi leads to an increase in the number of Ey^+^ (green) INPs (Dpn^+^ (red)). Mann-Whitney Test p=0.007, **; p<0.001, ***; p<0.001, ***. (**M**) Schematic summary of *scro* loss of function phenotype compared to wild type and *ey* loss of function. (**N**) Summary of the regulatory relationships between Hbn, *scro*, Ey and Grh. The grey arrows indicate previously established regulatory relationships. Single section confocal images unless stated otherwise. White dotted lines indicate *D*-GAL4>*mCD8-RFP* expression. Brains were dissected at wandering third instar stage. *n* = 6 brain lobes.

A further predicted regulatory relationship between temporal transcription factors is that each activates expression of the subsequent temporal factor and represses the next plus one. This would suggest that Hbn should activate Ey expression and repress Scro. In accordance with this prediction, we found that misexpression of Hbn in INPs activated the Ey temporal window precociously (Fig. 4C) and a reduction in *scro* (Fig. 4D). Consistent with these results, knockdown of *hbn* by RNAi (Fig. S4C) led to an increase in *scro* expression (Fig. 4E). We conclude that Hbn activates Ey and represses *scro* (Figs. 4G and 4H), the type of behaviour attributed to classically defined temporal transcription factors, exemplified in the embryonic temporal cascade (Isshiki et al., 2001).

### Hbn is sufficient to activate the middle-aged temporal factor Grh

We observed that the onset of Hbn expression coincided with the start of Grh expression. Furthermore, ectopic expression of Hbn resulted in concomitant expression of Grh and we noticed that INPs with stronger Hbn staining signal also correlated with higher Grh signal (Fig. S4D). As a result, we postulated that these two temporal factors shared a regulatory relationship. We found that ectopic Hbn expression was sufficient to activate Grh precociously and to extend its expression window in all DM lineages (Fig. S4E), even DM1, which normally does not express Grh. Grh was induced in almost all INPs in the DM1 lineage (Fig. 4F). Hbn is normally expressed at lower levels in DM1 than in other DMs, suggesting that Hbn may activate Grh in a dosage-dependent manner. Forced expression of Hbn in DM lineages drove INPs toward a “middle-aged fate”: most INPs remained D^-^ Grh^+^ Ey^+^ (Fig. 4G). We also observed that lineages DM1-4 and DM6 had significantly fewer INPs (Fig. S4E, F), suggesting that Hbn may regulate cellular longevity during the middle-aged window. Taken together, our results demonstrate that Hbn is a newly discovered temporal transcription factor that activates Grh and Ey expression and represses *scro* to progress INP temporal transitions (Fig. 4H).

### *scro* acts in a negative feedback loop with Ey

We found that *scro* expression was restricted to the oldest INPs, overlapping and extending beyond the Ey expression window (Fig. 3O and S3F). NanoDam revealed that Ey was bound at the *scro* locus, leading us to hypothesize that Ey might activate *scro* transcription in INPs. Ectopic expression of Ey was sufficient to activate *scro* precociously in all DM lineages except DM2 and 3, where a reduction in *scro* was observed (Fig. 4I and S4G). Conversely, knocking down *ey* expression in INPs lead to an almost complete loss of *scro* expression (Fig. 4J). We conclude that Ey is both necessary and sufficient to activate *scro* expression and that Ey is likely to act directly.

Given the reciprocal regulatory interactions observed between temporal factors, we tested whether *scro*, as the last identified temporal factor in INPs, represses Ey expression to terminate the Ey temporal window. In support of this hypothesis, we found that ectopic expression of Scro resulted in the loss of Ey (Fig. 4K). Next, we assessed whether the loss of *scro* in INPs would affect Ey expression. Using two independent *scro* RNAi constructs that knocked down expression effectively (Fig. S4H) we found that the loss of *scro* extended the Ey temporal window (Fig. 4L), without affecting the number of D^+^ or Grh^+^ INPs (Fig. S4I, J). Therefore, Scro is necessary and sufficient to close the Ey temporal expression window.

Interestingly, we observed that lineages lacking *scro* contained significantly more INPs than controls, suggesting an effect on longevity (Fig. S4K). It had been reported previously that the loss of Ey lead to an increased number of INPs (Bayraktar and Doe, 2013), comparable to the numbers we observed after the loss of *scro* (Fig. S4K). The effects on temporal patterning, however, were distinct. Removing Ey extended the middle-aged temporal window (INPs remained Grh^+^/Hbn^+^) (Fig. 4M) (Bayraktar and Doe, 2013). In contrast, in the absence of *scro* INPs progressed through the middle-aged window and, instead, the late temporal window was extended (INPs remained Ey^+^) (Fig. 4L and 4M). Together, our results indicate that *scro* is activated directly by Ey in INPs and that Scro restrains the Ey temporal window at the end of the temporal cascade (Fig. 4N).

### Hbn and Scro act as temporal factors in the developing visual system

Temporal patterning also regulates neuronal diversity in the developing optic lobe, where D and Ey are expressed temporally in NSCs in the medulla of the optic lobe (Fig. 5A) (Bayraktar and Doe, 2013; Li et al., 2013; Suzuki et al., 2013). Intriguingly, we found that the novel INP temporal factors we identified, Hbn and *scro*, were also expressed in subpopulations of medulla NSCs (Fig. 5B-C and Fig. S5A-B). As we had observed in INPs, we found that Hbn, Ey and *scro* were expressed sequentially in medulla NSCs (Fig. 5D-F). Interestingly, we could show that several of the regulatory interactions we discovered in INPs were conserved in medulla NSCs. We found that *scro* was necessary and sufficient for repression of Ey in NSCs (Fig. 5G-H), indicating that termination of the Ey temporal window by *scro* is conserved between INPs and the optic lobe. In addition, misexpression of Hbn dramatically reduced *scro* expression, fulfilling the ‘repression of next plus one’ rule (Fig. 5I). A comparison of Ey and D binding in INPs and the optic lobe revealed slightly different patterns at the *hbn* and *scro* loci (Fig. 5J-K): D binds strongly at the *hbn* locus in INPs, but more weakly in the optic lobe. This may reflect differential regulation in the two cell types. For example, in contrast to INPs, in the optic lobe D is expressed after Hbn and the D and Hbn temporal windows do not overlap (Fig. S5C). We conclude that the temporal expression cascade, from Hbn to Ey to *scro*, is conserved in progenitors that generate both the central complex of the brain and the visual processing system (Fig. 5L-M). This is particularly striking as INPs and optic lobe NSCs are distinct progenitors with different origins, the former born from asymmetrically dividing Type II NSCs and the latter derived from symmetrically dividing neuroepithelial cells (Egger et al., 2007).

**Figure 5:**
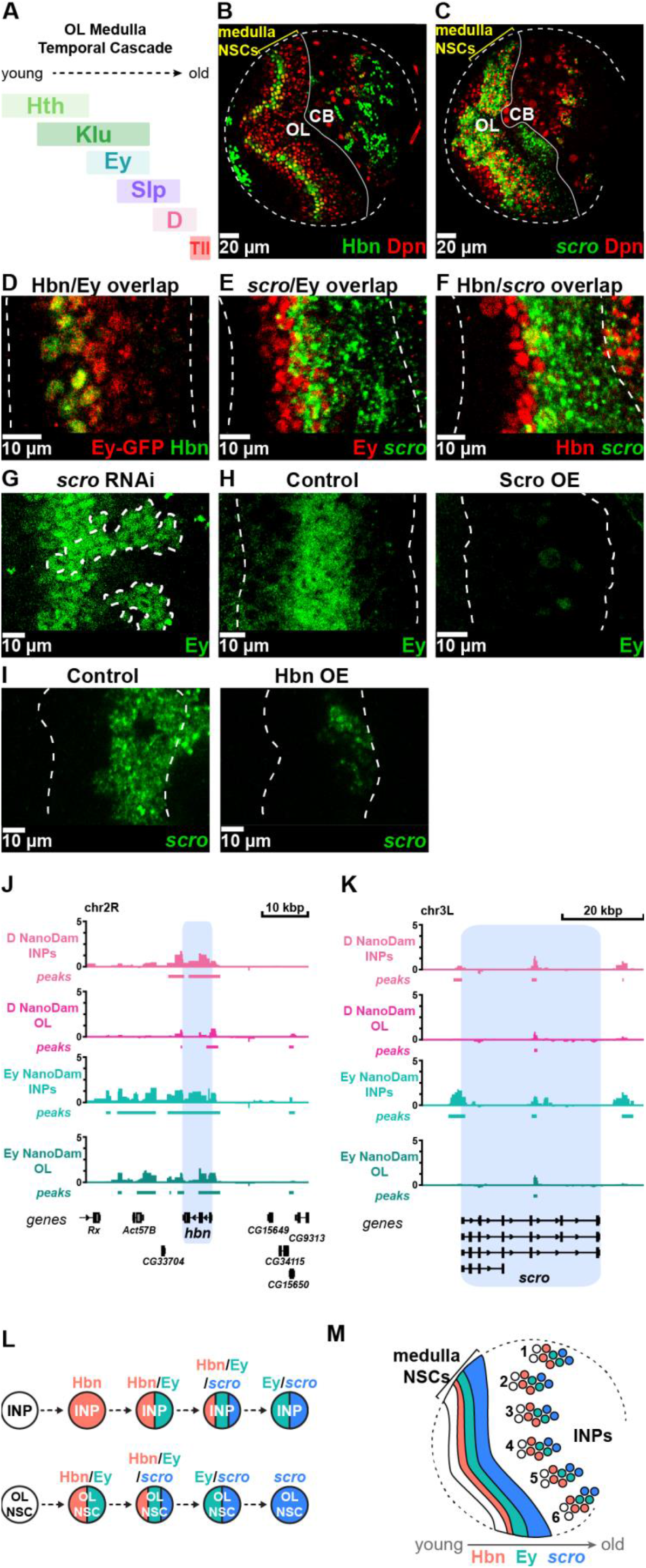
Hbn and *scro* are expressed in temporal windows in the optic lobe temporal cascade. (**A**) Schematic showing the temporal cascade of the optic lobe (OL) medulla NSCs. (**B**) Hbn (green) is expressed in the OL medulla NSCs (Dpn^+^ (red) and highlighted with yellow bracket) in addition to the central brain (CB). (**C**) *scro* (green) is expressed in the OL medulla NSCs (Dpn^+^ (red) and highlighted with yellow bracket) in addition to the CB. (**D**) Hbn (green) expression coincides with the start of the Ey temporal window (Ey-GFP^+^, (red)) in medulla NSCs. (**E**) *scro* (green) expression begins in the second half of the Ey window (Ey, red). (**F**) *scro* (green) is expressed in the oldest Hbn^+^ (red) OL medulla NSCs. (**G**) Loss of *scro* extends Ey expression (green) in medulla NSCs. Clones expressing *scro* RNAi^TRiP^ are indicated with white dotted outlines. (**H**) Ectopic Scro expression in OL NSCs results in the loss of the Ey temporal window (green). *insc*-GAL4 was used to drive UAS-*scro*. (**I**) Ectopic Hbn expression in OL NSCs leads to a reduction of s*cro* expression (green). *insc*-GAL4 was used to drive UAS-*hbn*. (**J**) NanoDam binding profiles of D and Ey across the *hbn* gene locus (blue) in INPs and OL NSCs. *ogre*-GAL4 was used to drive NanoDam in OL NSCS in combination with D-GFP or Ey-GFP. (**K**) NanoDam binding profiles of D and Ey across the *scro* gene locus (blue) in INPs and OL NSCs. (**L**) The order of Hbn, Ey and *scro* expression is conserved in the INP and OL medulla NSC temporal cascades. Single section confocal images. Dotted white lines in B and B outline the edges of the brain lobes; in D-I indicate the region containing medulla NSCs. (**M**) Schematic showing the conserved expression of Hbn, Ey and *scro* in the medulla NSCs and INPs of the DM Type II lineages (1-6). See also Figure S5.

## Discussion

Temporal patterning leads to the generation of neuronal diversity from a relatively small pool of neural stem or progenitor cells. Temporal regulation is achieved by the restricted expression of temporal transcription factors within precise developmental windows. The onset and duration of each temporal window in neural stem or progenitor cells must be regulated tightly in order for the appropriate subtypes of neurons to be generated at the correct time to establish functional neuronal circuits.

Here we focused on the INPs of the Type II NSC lineages that generate the central complex of the *Drosophila* brain. Previously, INPs were shown to express sequentially the temporal factors D, Grh and Ey (Bayraktar and Doe, 2013). Given the expectation that other temporal factors remained to be discovered, and that these were likely to be the transcriptional targets of the known temporal factors, we developed a new technique called NanoDam to profile the binding targets of D, Grh and Ey with cell-type specificity and within their individual temporal windows.

NanoDam enables genome-wide profiling of any endogenously tagged chromatin-binding protein with a simple genetic cross, bypassing the need to generate Dam-fusion proteins, or the need for specific antisera or cell isolation. Furthermore, NanoDam profiles binding only in cells where the tagged protein is normally expressed. Binding within a subset of the protein’s expression pattern can be achieved by controlling NanoDam with specific GAL4 drivers (Fig. 1D). To date, collaborative efforts have produced more than 3900 *Drosophila* lines expressing GFP-tagged proteins in their endogenous patterns (Table S2). Approximately 93% of all transcription factors have been GFP-tagged in lines that are publicly available at stock centres. Lines that are not yet available can be rapidly generated by CRISPR/Cas9-mediated tagging. NanoDam is thus a versatile tool that can be used as a higher throughput method to profile genome-wide binding sites of any chromatin associated protein. NanoDam can be readily adapted for use in other organisms to facilitate simpler and easier *in vivo* profiling experiments, as we have demonstrated previously for TaDa (Cheetham et al., 2018; Aloia et al., 2019).

By combining the power of NanoDam with scRNA-seq, we were able to identify *scro* and *hbn* as novel temporal factors in the INPs of Type II NSC lineages. We showed that *hbn* and *scro* regulate the maintenance and transition of the middle-aged and late temporal windows. The mammalian homologues of *ey* (*Pax6*), *hbn* (*Arx*) and *scro* (*Nkx2*.*1*) are restricted to distinct progenitor populations in the developing mouse forebrain (Colombo et al., 2004; Götz et al., 1998; Sussel et al., 1999). We found that *scro* regulates the late INP identity by repression of Ey. Interestingly, the loss of *Nkx2*.*1* in the mouse forebrain leads to aberrant expression in ventral regions of the dorsal factor *Pax6* (Sussel et al., 1999), suggesting that the repressive relationship between *scro* and *ey* may be conserved between *Nkx2*.*1* and *Pax6*. Not all relationships appear to be conserved, however. We found that Hbn promotes progression through the middle-aged temporal stage and that maintenance of the middle-aged temporal window is regulated in part by interactions between Hbn and Grh. *Arx* mutant mice exhibit loss of upper layer (later-born) neurons but no change in the number of lower layer (early-born) neurons (Colasante et al., 2015).

Intriguingly, the novel temporal factors identified in the INPs were also temporally expressed in optic lobe NSCs (Fig. 5L) and the regulatory relationships between *scro* and Ey appeared to be conserved. This suggests that similar regulatory strategies may be shared between neural stem cells or progenitor cells in order to regulate longevity and neuronal subtype production. The remarkable conservation of the regulatory interactions of *scro* in two different progenitor cell types with different origins in the *Drosophila* brain may also be translated to the context of mammalian neurogenesis, highlighting the possibility of a more generalised regulatory network used by stem and progenitor cells to regulate cell fate, progeny fate and proliferation.

The Type II lineages in *Drosophila* divide in a very similar manner to the outer radial glia (oRGs) that have been attributed to the rapid evolutionary expansion of the neocortex seen in humans and other mammals (Fietz et al., 2010; García-Moreno et al., 2012; Hansen et al., 2010; Wang et al., 2011). Interestingly, oRGs show a shortened cell cycle length in primates (Penisson et al., 2019) in comparison to rodent progenitors, which increase cell cycle duration as development progresses (Betizeau et al., 2013). Investigating whether oRGs use temporally expressed factors to control longevity and cell cycle dynamics at different developmental stages in order to regulate neuronal subtype generation would be important for understanding neocortex development.

There is significant heterogeneity between the Type II lineages and our study has identified differences in the regulatory relationships of *hbn* and *scro*. For example, misexpression of Ey leads to an increase in *scro* in all lineages except DM 2 and 3, where *scro* expression is reduced. To date, Hbn is the only factor identified that activates Grh in DM1, the lineage that does not normally express Grh. The heterogeneity between lineages may be a consequence of variations in combinatorial binding of temporal factors, as our NanoDam data indicate. Although INPs share temporal factors, different DM lineages display subtle to striking differences when the temporal cascade is manipulated, demonstrating the likelihood that each DM employs unique temporal cascades. Combinatorial binding would enable more complex regulatory interactions that could refine or sub-divide temporal windows in the INPs.

## Acknowledgements

We thank Markus Affolter, Michael Akam, Alex Gould, Stephan Thor, Uwe Walldorf, Bloomington *Drosophila* Stock Center and the Vienna *Drosophila* Resource Center (VDRC) for reagents. We acknowledge Katarzyna Kania at the Cancer Research UK Cambridge Institute Genomics Core Facility for performing sample preparation and sequencing of 10X Chromium samples.

This work was funded by Wellcome Trust PhD Studentships to E.G.C (089616), A.E.H. (102454) and J.L.Y.T. (203798), an EMBO Long Term Fellowship to P.M.F. and Wellcome Trust Programme Grant (092545), Wellcome Trust Senior Investigator Award (103792) and Royal Society Darwin Trust Research Professorship to A.H.B. A.H.B acknowledges core funding to the Gurdon Institute from the Wellcome Trust (092096) and CRUK (C6946/A14492).

## Author contributions

A.E.H., P.M.F. and A.H.B. conceptualised the project. A.E.H., J.L.Y.T., E.G.C. and P.M.F. performed the experiments. T.S. and R.K. assisted with generation of single-cell RNA-seq and NanoDam datasets. R.K. analysed single-cell RNA-seq and NanoDam binding data. A.E.H., J.L.Y.T and A.H.B. wrote the manuscript. All authors reviewed and commented on the manuscript.

## Methods

### Fly Stocks

*Drosophila melanogaster* were reared in cages at 25 °C. Embryos were collected on yeasted apple juice plates. For experiments involving GAL80^ts^ embryos were kept at 18 °C until hatching. After hatching, larvae were transferred to a yeasted food plate and reared to wandering third larval instar stage before dissection.

The following lines were used to drive transgenes under the control of UAS in a spatially and temporally restricted manner: *D*-GAL4 (GMR12E09-GAL4, BDSC 48510), *insc*-GAL4 (*GAL4*^*MZ1407*^) (Luo et al., 1994), *ogre*-GAL4 (GMR29C07-GAL4, BDSC 49340), *wor*-GAL4 (Albertson et al., 2004), *tub*-GAL80^ts^ (BDSC 7019), Ay-GAL4, UAS-GFP (BDSC 4411), 10xUAS-*IVS-mCD8-RFP* (BDSC 32219), UAS-mCD8-GFP (BDSC 5130), UAS-*Dicer2* (VDRC 60008) and *hsFLP*^122^. The fusion proteins used were: *cph*::YFP (CPTI-001740, PMF and AHB, in preparation), D-GFP (BDSC 66758), Grh-GFP (BDSC 42272), Ey-GFP (BDSC 42271). The misexpression lines used were: UAS-*D* (BDSC 8861), UAS-*ey* (BDSC 6294), UAS-*grh* (BDSC 42227), UAS-*hbn* (this study), UAS-*scro* (this study). The RNAi lines used were: UAS-*ey*-RNAi (VDRC 106628), UAS-*grh*-RNAi (VDRC 101428), UAS-*hbn*-RNAi^CDS2^ (this study), UAS-*hbn*-RNAi^3UTR^ (this study), UAS-*mCherry*-RNAi (BDSC 35785), UAS-*scro*-RNAi (BL33890) and UAS-*scro*-RNAi^CDS^ (this study). The generation of UAS-*NanoDam* is described below. *w*^*1118*^ was used for NanoDam control experiments and as a reference stock for functional experiments.

### Generation of expression constructs

pUAST-mCherry-Dam-NLS-vhhGFP4 (pUAST-NanoDam) was generated by PCR amplifying vhhGFP4 from genomic DNA isolated from deGradFP flies (Caussinus et al., 2012) using the primers Forward: AGTGGCGGTGGGCCAAAAAAGAAAAGAAAAAGTCCGCGGCAAATGGATCAAGTCCAACTGGTGG and Reverse: ATGTCACACCACAGAAGTAAGGTTCCTTCACAAAGATCCTTAGCTGGAGACGGT GACCTG. The resulting PCR product was cloned into the pUAST-mCherry-Dam vector (Southall et al., 2013) with XhoI and XbaI sites using Gibson Assembly.

An NLS sequence (AGATCTCATGCCAAAAAAGAAAAGAAAAAGTGGCGGTGGGCCAAAAAAGAAAAGAAAAAGTCCGCGGCCGC, generated by annealing complementary oligos) was inserted between the BglII and SacII cloning sites.

pUAST-attB-hbn was generated by PCR amplifying the *hbn* coding sequence from an embryonic cDNA library using the primers Forward: AGATGAATTCATGATGACCACGACGACCTCG and Reverse: ATGACTCGAGTCAGTCCTCGCCCTTGGTG. The resulting PCR product was cloned into the pUAST-attB vector (Bischof et al., 2007) with EcoRI and XhoI sites.

pUAST-attB-scro was generated by PCR amplifying the *scro-RA* coding sequence from an embryonic cDNA library using the primers Forward: TACCAGGAATTCATGTCATCGCACGGCCTTGCTTAC and Reverse: TAGTATGCGGCCGCTTACCATGCCCGGCCTTGTAAGGG. The resulting PCR product was cloned into the pUAST-attB vector (Bischof et al., 2007) with EcoRI and NotI sites.

### Short hairpin RNAi generation

Short hairpin *scro* RNAi constructs, pWALIUM20-*scro*-shRNA^5UTR^ (targets *scro* 5UTR) and pWALIUM20*-scro*-shRNA^CDS^ (targets *scro* CDS), were generated by annealing 10 µM concentration of the following oligonucleotide pairs at 98 °C for 5 minutes in annealing buffer (10 mM Tris pH 7.5, 0.1 M NaCl, 1 mM EDTA) and then leaving the mixture to cool at room temperature. The resulting products were cloned into the pWALIUM20 vector (Perkins et al., 2015) with EcoRI and Nhe1 sites.

*hbn*-shRNA^CDS2^ Forward ctagcagtGTTCGAAGATCTCGCACAACAtagttatattcaagcataTGTTGTGCGAGATCTTCGAACgcg and *hbn*-shRNA^CDS2^ Reverse aattcgcGTTCGAAGATCTCGCACAACAtatgcttgaatataactaTGTTGTGCGAGATCTTCGAACactg; *hbn*-shRNA^3UTR^ Forward ctagcagtGATTCGTTATGTACATATATAtagttatattcaagcataTATATATGTACATAACGAATCgcg and *hbn*-shRNA^3UTR^ Reverse aattcgcGATTCGTTATGTACATATATAtatgcttgaatataactaTATATATGTACATAACGAATCactg; *scro*-shRNA^5UTR^ Forward ctagcagtAGTGGATATTCATAATAAATTtagttatattcaagcataAATTTATTATGAATATCCACTgcg and *scro*-shRNA^5UTR^ Reverse aattcgcAGTGGATATTCATAATAAATTtatgcttgaatataacta AATTTATTATGAATATCCACTactg; *scro*-shRNA^CDS^ Forward ctagcagtGCCCAGGTGTACAGACCTATTtagttatattcaagcataAATAGGTCTGTACACCTGGGCgcg and *scro*-shRNA^CDS^ Reverse aattcgcGCCCAGGTGTACAGACCTATTtatgcttgaatataactaAATAGGTCTGTACACCTGGGCactg. For all expression and RNAi constructs, transgenic flies were generated by germline injection into embryos carrying *y, v, nos*-phiC integrase; *attP40*; + (BDSC 25709).

### NanoDam Experimental Design

To perform Cph NanoDam, a Cph::YFP; UAS-*NanoDam* line was crossed to Cph::YFP; *wor*-GAL4. As a control, UAS-*NanoDam* was crossed to *wor*-GAL4. *wor*-GAL4 is expressed in neuroblasts from approximately stage 10. Embryos were collected at 25 °C and harvested at stage 16. 10-15 µl of embryos were harvested for DNA purification.

For INP NanoDam, flies carrying UAS-*NanoDam, tub*-GAL80^ts^ and *D*-GAL4 were crossed to *w*^*1118*^ (control), D-GFP, Grh-GFP or Ey-GFP. For optic lobe NanoDam, flies carrying UAS-*NanoDam, tub*-GAL80^ts^ and *ogre*-GAL4 were crossed to *w*^*1118*^, D-GFP, or Ey-GFP. Temporal restriction of NanoDam expression was achieved using GAL80^ts^, a temperature-sensitive negative regulator of GAL4 (Matsumoto et al., 1978; McGuire et al., 2003). Embryos were collected on yeasted apple juice plates at 25 °C and then transferred at 18 °C. Newly hatched larvae were transferred to yeasted food plates and raised at 18 °C for 6 days before shifting to 29 °C for 14 hours. Brains were dissected from wandering third instar larvae in PBS and then transferred to ice-cold PBS. Genomic DNA was extracted from approximately 50 brains per sample.

### NanoDam sample processing

NanoDam samples were processed using the DamID-seq protocol as described previously (Marshall et al., 2016). Sequencing was performed as single end 50 bp reads generated by an Illumina HiSeq 1500 at the Gurdon Institute NGS Core Facility. Details of biological replicates performed can be found in Figure S1.

### NanoDam data processing

The quality of all *.fastq-files was validated by using FastQC (v0.11.5). Data processing was performed with a wrapper script to automate and parallelize the application of the damidseq_pipeline (Marshall and Brand, 2015) with the slurm workload manager (v15.08.13). All *.fastq.gz-files were mapped with bowtie2 (v2.3.4.1) to the *Drosophila* dm6 genome assembly and all reads were assigned to bins defined by consecutive GATC sites throughout the genome. The NanoDam profile of Cph was normalised against a control sample that expressed the NanoDam transgene in the absence of YFP. For NanoDam of each transcription factor, all replicates (4 D-GFP replicates, 5 Grh-GFP replicates, and 4 Ey-GFP replicates) were normalised individually to all control replicates (8 *w*^*1118*^ replicates) followed by quantile normalisation of pairwise comparisons to each other. Binding profiles for each transcription factor were generated by averaging the binding intensities across all normalised comparisons per GATC-bin. The averaged logarithmic binding intensities were subsequently backtransformed and bedGraphToBigWig (v4) was used to generate *.bw-files for visualisation of the binding profiles in IGV (Robinson et al., 2011). Broad peaks were called with macs2 (v2.1.2.) on *.bam-files derived from the damidseq_pipeline for all NanoDam/control pairwise combinations. Overlapping peak regions were merged with bedtools (v2.26.0) into consensus peaks. Consensus peaks were filtered by false discovery rate (i.e., FDR<10^−25^) and by occurrence in more than 50 % across all pairwise combinations.

Binding intensities for all GATC-bins overlapping with individual peaks were averaged and normalised for the length of the peak. Intensities for GATC-bins intersecting with the peak borders were weighted depending on the overlap of the respective bin with the peak. The resulting intensities per peak and genotype were converted into z-scores and clustered with the ‘kmeans’ function of the R ‘stats’ package (v3.6.1). The optimal number of clusters was determined by using the factoextra (v1.0.5), clValid (v0.6-6) and mclust (v5.4.5) packages in R. All considered clustering approaches were evaluated by calculating silhouettes with the cluster (v2.0.7-1) package.

Peaks were assigned to the closest transcriptional start site of protein-coding genes according to ensembl annotations with bedtools (dm6, biomaRt v2.38.0). To identify genes encoding transcription factors the resulting genes were intersected with the curated list of supported *Drosophila* transcription factors from FlyTF.org (https://www.mrc-lmb.cam.ac.uk/genomes/FlyTF/old_index.html).

### Single cell sequencing sample preparation

Embryos of the genotypes *w*; 10xUAS-*IVS-mCD8-RFP*; *D*-GAL4 or *w*; 10xUAS-*IVS-mCD8-RFP*/D-GFP; *D*-GAL4/*+* were collected on yeasted apple juice plates at 25 °C. Newly hatched larvae were transferred to yeasted food plates and reared at 25 °C until wandering third instar stage. Sample preparation prior to FACS was performed as (Harzer et al., 2013), but with the use of PBS in place of Schneider’s medium. In brief, larvae were washed in 70 % EtOH/PBS for 1 minute before dissection. Brains from each genotype were then dissected in PBS for one replicate each, transferred to 1.5 ml low-binding tubes with ice-cold Rinaldini solution and then rinsed twice with ice-cold Rinaldini solution. Brains were incubated for 1 hour at 30 °C in dissociation solution (Schneider’s Insect Medium with 1 mg/ml Collagenase I and 1 mg/ml Papain) then rinsed with ice-cold Rinaldini solution, followed by sterile filtered 0.03 % BSA/PBS. The solution was pipetted up and down to dissociate the brains and the resulting cell suspension was passed through a 10 µm mesh filter into a 5 ml FACS tube. SYTOX Blue Dead Cell Stain (Invitrogen) was added to the cell suspension before proceeding to FACS.

FACS was performed using the SH800Z Cell Sorter (Sony) at 4 °C. Single cells were sorted into PBS with 0.03 % Bovine Serum Albumin to prevent clumping. Live, single cells were sorted based on size, and fluorescence intensities of RFP and SYTOX Blue Dead Cell Stain. Gates for FACS were established using a negative control to adjust for autofluorescence and a positive control for SYTOX Blue Dead Cell Stain using dissociated cells incubated at 65 °C for 15 minutes.

Single cell RNA sequencing libraries were generated using 10X Chromium (v2 chemistry kit, 10X Genomics) and sequenced on an Illumina HiSeq 4000 at the Genomics Core Facility, Cancer Research UK Cambridge Institute.

### Single cell sequencing data processing

The acquired single cell RNA sequencing files were mapped and count matrices derived from Cell Ranger (v2.2.1). The required reference transcriptome was build from *.gtf- and *.fa-files for the Ensembl dm6 genome assembly. The Dichaete-BAC used to generate the D-GFP (BL66758) was in silico cloned and custom made *.gtf- and *.fa-files of the resulting plasmid were combined with the *Drosophila* transcriptome to generate a separate reference for validation of D-GFP expression as a means to quality check the sample preparation. Replicates (with and without D-GFP) were found to be very similar when the two samples were integrated. Data for both single cell data sets were separately normalized and scaled with the Seurat R package (v2.3.4) prior to their integration via canonical correlation analysis (Butler et al., 2018; Stuart et al., 2019). The number of screened and chosen genes (i.e., ‘num.possible.genes’ and ‘num.genes’) while constructing the metagene as well as the number of aligned dimensions were optimized during the subspace alignment. After excluding tracheal clusters (i.e., 5 and 6), significantly differentially expressed genes for all clusters were identified by using the ‘FindMarkers’ command (i.e., “Wilcoxon rank sum test”). To identify transcription factor genes, the list was intersected with the aforementioned list of supported *Drosophila* transcription factors from FlyTF.org.

### Sample fixation and immunostaining

Larval brains were dissected in PBS and fixed on a shaker for 20 minutes in 4 % formaldehyde/PBS. Fixed brains were washed well with PBS containing 0.3 % Triton-X (PBTx) before immunostaining and then for at least 15 minutes in 10 % normal goat serum/PBS. Samples were incubated overnight at 4 °C with primary antibodies diluted in 0.3 % PBTx, washed well with 0.3 % PBTx, then incubated overnight at 4 °C with secondary antibodies diluted in 0.3 % PBTx. Samples were washed well with 0.3 % PBTx then mounted in Vectashield (Vector laboratories) for imaging.

The following primary antisera were used: guinea pig anti-D 1:200 (a gift from Alex Gould), guinea pig anti-Dpn 1:5,000 (Caygill and Brand, 2017), rat anti-Dpn 1:100 (abcam, 11D1BC7, ab195173), rabbit anti-Ey (1:300) (a gift from Uwe Walldorf), chicken anti-GFP 1:2,000 (abcam, ab13970), rat anti-Grh 1:1000 (Baumgardt et al., 2009), rabbit anti-Hbn (1:200) (a gift from Uwe Walldorf), guinea pig anti-Opa 1:500 (a gift from Michael Akam). Secondary antibodies conjugated Alexa Fluor dyes (Life Technologies) or DyLight-405 1:200 (Jackson Laboratories) were used to detect primary antibodies.

### *In situ* hybridisation chain reaction (HCR)

To perform HCR for *scro* mRNA, custom probes designed against the *scro* coding sequence, buffers and fluorophore-labelled amplification hairpins were sourced from Molecular Instruments (Choi et al., 2018). The HCR protocol was adapted for use on third instar *Drosophila* larval brains (Choi et al., 2016). In brief, larval brains were fixed on a shaker at room temperature for 20 minutes in 4 % formaldehyde/PBS. Fixed brains were washed well with PBS and then incubated in probe hybridisation buffer (PHB) (Molecular Instruments) at 37 °C for 10 minutes. PHB was removed from the samples and replaced with pre-warmed probe mix (0.8 µl of probe added to 200 µl PHB). Samples were incubated in probe mix overnight at 37 °C. The following day, samples were washed well with probe wash buffer (PWB) (Molecular Instruments) at 37 °C and then washed with 5XSSC containing 0.1 % Triton-X (5X SSCTx) at room temperature. Samples were incubated in amplification buffer (AB) (Molecular Instruments) at room temperature for at least 10 minutes. AB was removed and replaced with 100 µl AB containing 2 µl each fluorophore-labelled amplification hairpin1 and hairpin2 (Molecular Instruments). Samples were then incubated overnight in the dark at room temperature. Note that hairpins 1 and 2 were heated separately at 95 °C for 1.5 minutes and cooled at room temperature for 30 minutes in the dark before use. The next day, samples were washed well with 5X SSCTx before proceeding to immunostaining processing for antibody co-staining. Amplification hairpins were labelled with Alexa Fluor 488 or 647. HCR samples were mounted in SlowFade Gold antifade reagent (Invitrogen) for imaging.

### Image acquisition and analysis

Fluorescent images were acquired using a Leica SP8 confocal microscope. Images were analyses using Fiji (Schindelin et al., 2012), which was also used to adjust brightness and contrast in images. Adobe Illustrator was used to compile figures. GraphPad Prism 8 for Mac OS X (www.graphpad.com) was used for statistical analyses. Following normality tests, Mann-Whitney U tests were used to assess the statistical significance between two genotypes and Kruskal-Wallis tests were used when experiments contained more than two genotypes.

## Tables

**Table S1.** Genes bound by D/Grh/Ey (NanoDam) and expressed in INPs (scRNA-seq).

| ND1 | ND2 | ND3 | ND4 | ND5 | ND6 |
| --- | --- | --- | --- | --- | --- |
| *apt* | *ase* | *apt* | *Alh* | *Alh* | *BEAF-32* |
| *ftz-f1* | *cas* | *E2f1* | *apt* | *BEAF-32* | *E2f2* |
| *grh* | *dpn* | *Hr4* | *CrebA* | *E2f1* | *fd102C* |
| *ham* | *ftz-f1* | *ktub* | *D* | *ey* | *ftz-f1* |
| *hng3* | *grh* | *Optix* | *E2f1* | *fd102C* | *grh* |
| *klu* | *jumu* | *ph-p* | *ey* | *ham* | *sna* |
| *Myb* | *ken* | *sna* | *ham* | *hng3* | *SoxN* |
| *slp2* | *opa* | *SoxN* | *hbn* | *opa* |  |
| *sna* | *Optix* | *su(Hw)* | *klu* | *run* |  |
| *SoxN* | *slp2* | *Su(var)2-10* | *opa* | *scro* |  |
|  |  |  | *Optix* |  |  |
|  |  |  | *ph-p* |  |  |
|  |  |  | *run* |  |  |
|  |  |  | *slp2* |  |  |
|  |  |  | *SoxN* |  |  |
|  |  |  | *tap* |  |  |
|  |  |  | *wor* |  |  |

**Table S2.**
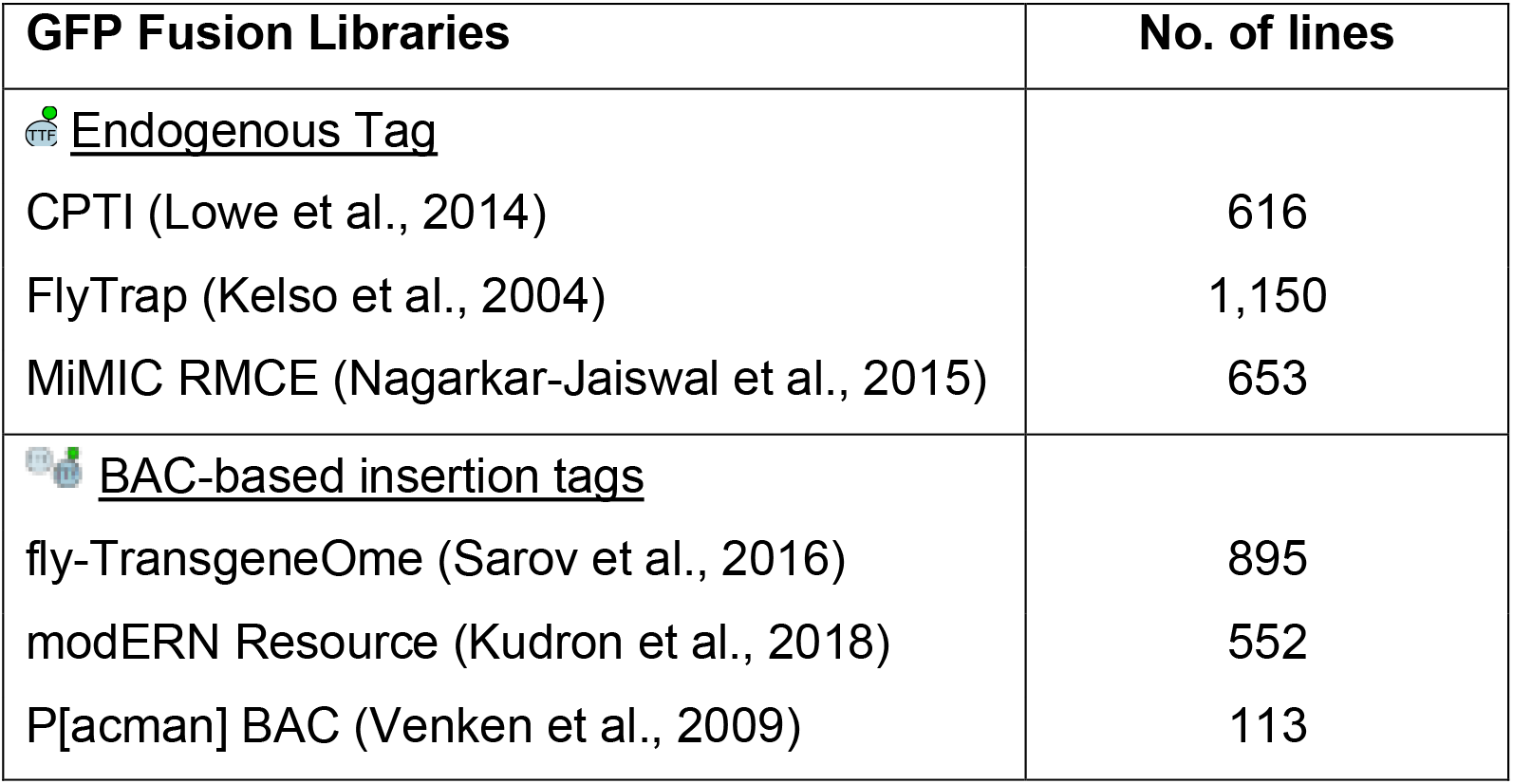
GFP-fusion libraries publicly available.

