## Supplemental Figures for "NanoDam identifies novel temporal transcription factors conserved between the *Drosophila* central brain and visual system" (from another version of this preprint: version 1, 2021-06-07)

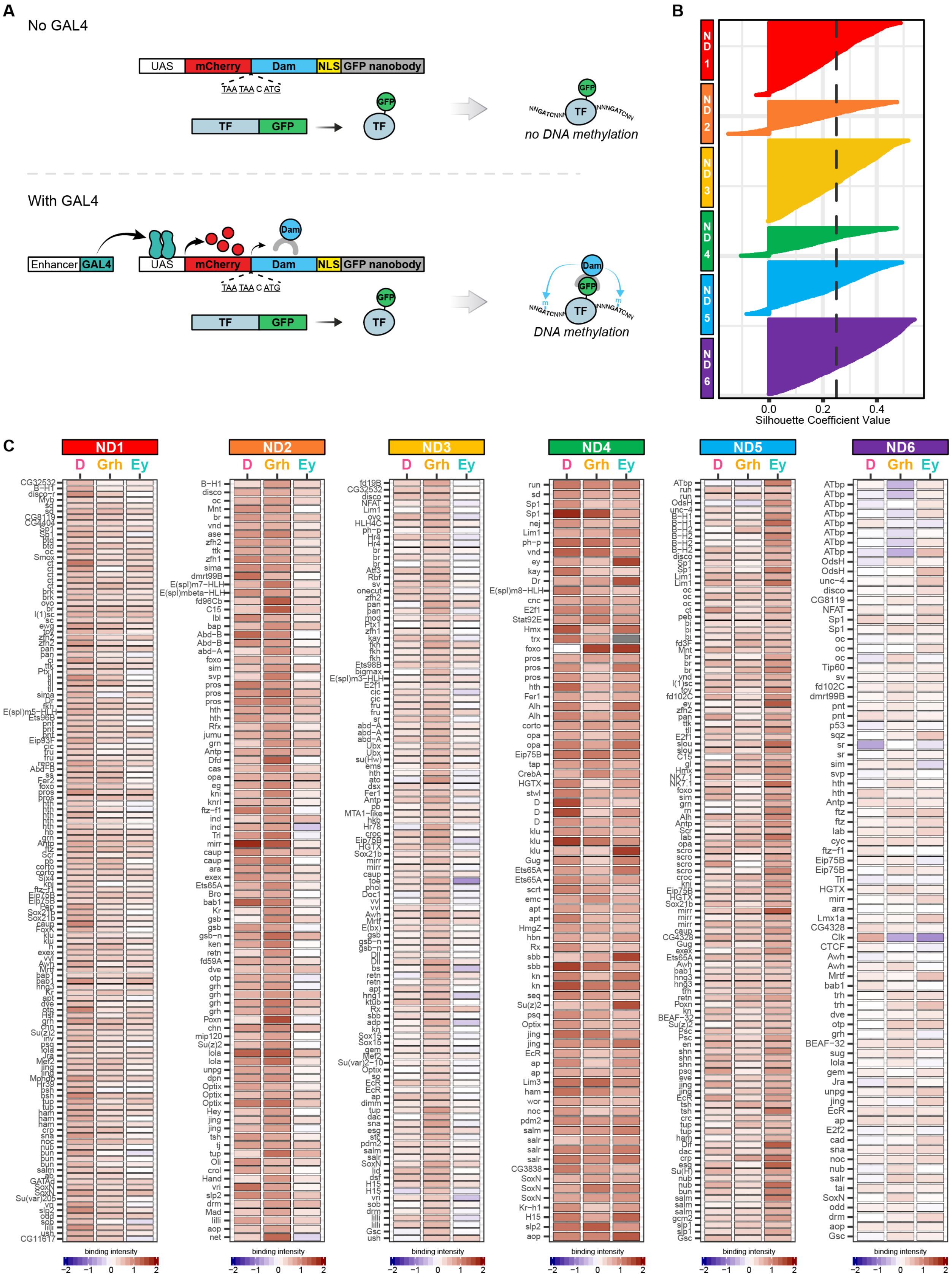
A
No GAL4
UAS
mCherry
Dam
NLS
GFP nanobody
TAA TAA C ATG
TF
GFP
no DNA methylation
With GAL4
Enhancer
GAL4
DNA methylation
B
ND1
ND2
ND3
ND4
ND5
ND6
0.0
0.2
0.4
Silhouette Coefficient Value
C
D
Grh
Ey
binding intensity
−2
−1
0
1
2

**Figure S1: NanoDam methodology.**

(**A**) Schematic representation of the NanoDam method. The NanoDam construct (the GFP nanobody fused to the C-terminus of Dam methylase with an intervening nuclear localisation sequence (NLS)) is under the control of UAS, allowing its expression to be activated in a cell-type-specific manner by GAL4. Translation of the NanoDam construct is greatly reduced by the addition of an upstream ORF (mCherry) followed by two stop codons, which prevents NanoDam expression in uninduced cells and nonspecific methylation (caused by high NanoDam levels) in induced cells.

(**B**) Average binding intensities for D, Grh and Ey across all peaks per NanoDam cluster represented as box plots. Silhouette analysis for all peaks provides an indicator for the robustness of peak assignment to its respective cluster. Dashed line represents the average silhouette length across all clusters.

(**C**) For each cluster (ND1-6), peaks were assigned to genes and then subsetted for transcription factors to generate the lists shown. NanoDam binding intensities for D, Grh or Ey are shown as z-scores for individual peaks. Each box represents the intensity of a single peak.

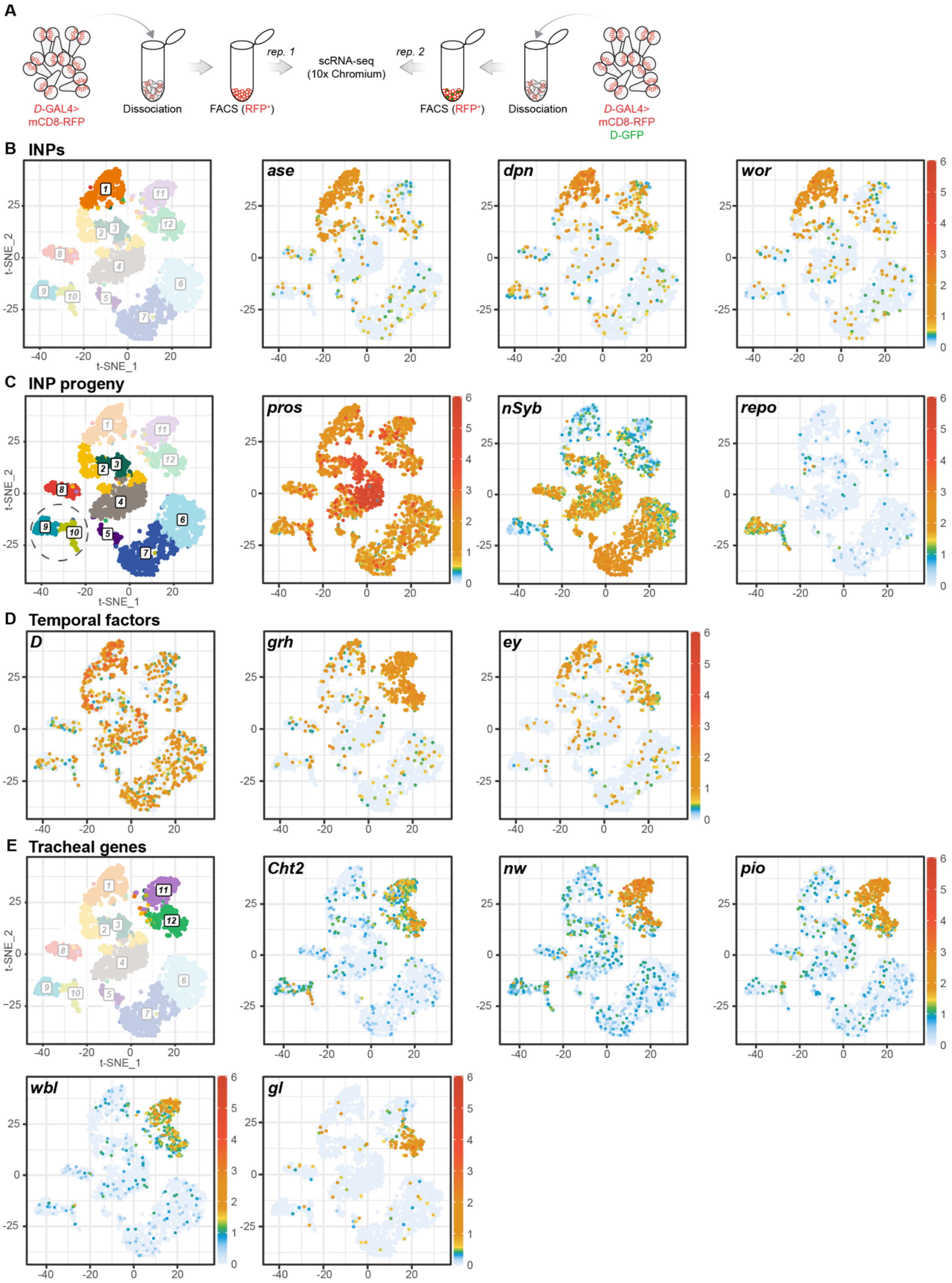
A
D-GAL4>
mCD8-RFP
Dissociation
FACS (RFP+)
rep. 1
scRNA-seq
(10x Chromium)
rep. 2
FACS (RFP+)
Dissociation
D-GAL4>
mCD8-RFP
D-GFP
B INPs
ase
dpn
wor
t-SNE_1
t-SNE_2
C INP progeny
pros
nSyb
repo
D Temporal factors
D
grh
ey
E Tracheal genes
Cht2
nw
pio
wbl
gl

**Figure S2: Assigning cell identities using known marker genes.**

(**A**) Schematic overview of the single cell RNA sequencing (scRNA-seq) approach. We performed two replicates: replicate 1 (rep.1) was performed on brains carrying *D*-GAL4>mCD8-RFP (right) and replicate 2 (rep. 2) on brains carrying *D*-GAL4>mCD8-RFP and D-GFP (left). Brains were dissected at wandering third instar stage, dissociated and then cells were sorted based on RFP expression before being submitted for 10x Chromium scRNA-seq.

(**B**) t-SNE visualisation of 4,086 cells coloured by cluster assignment highlighting Cluster 1, which corresponds to INPs. Cluster 1 was designated as INPs due to enriched expression of *asense* (*ase*), *deadpan* (*dpn*), *worniu* (*wor*).

(**C**) Clusters 2-10 contain a mixture of INP progeny and express high levels of *prospero* (*pros*). Clusters 2-8 are express *neuronal Synaptobrevin* (*nSyb*) and so are classified as neurons. Clusters 9 and 10 (outline with dotted circle) correspond to glia due to high expression of *reversed polarity* (*repo*).

(**D**) t-SNE plots coloured by expression of the temporal factors in INPs: *Dichaete (D)*, *grainyhead* (*grh*) and *eyeless (ey).*

(**E**) Clusters 11 and 12 correspond to presumed trachea due to strong enrichment of *Chitinase 2* (*Cht2*), *narrow* (*nw*) *piopio* (*pio*) (Affolter and Caussinus, 2008), *windbeutel* (*wbl*), *glass* (*gl*).

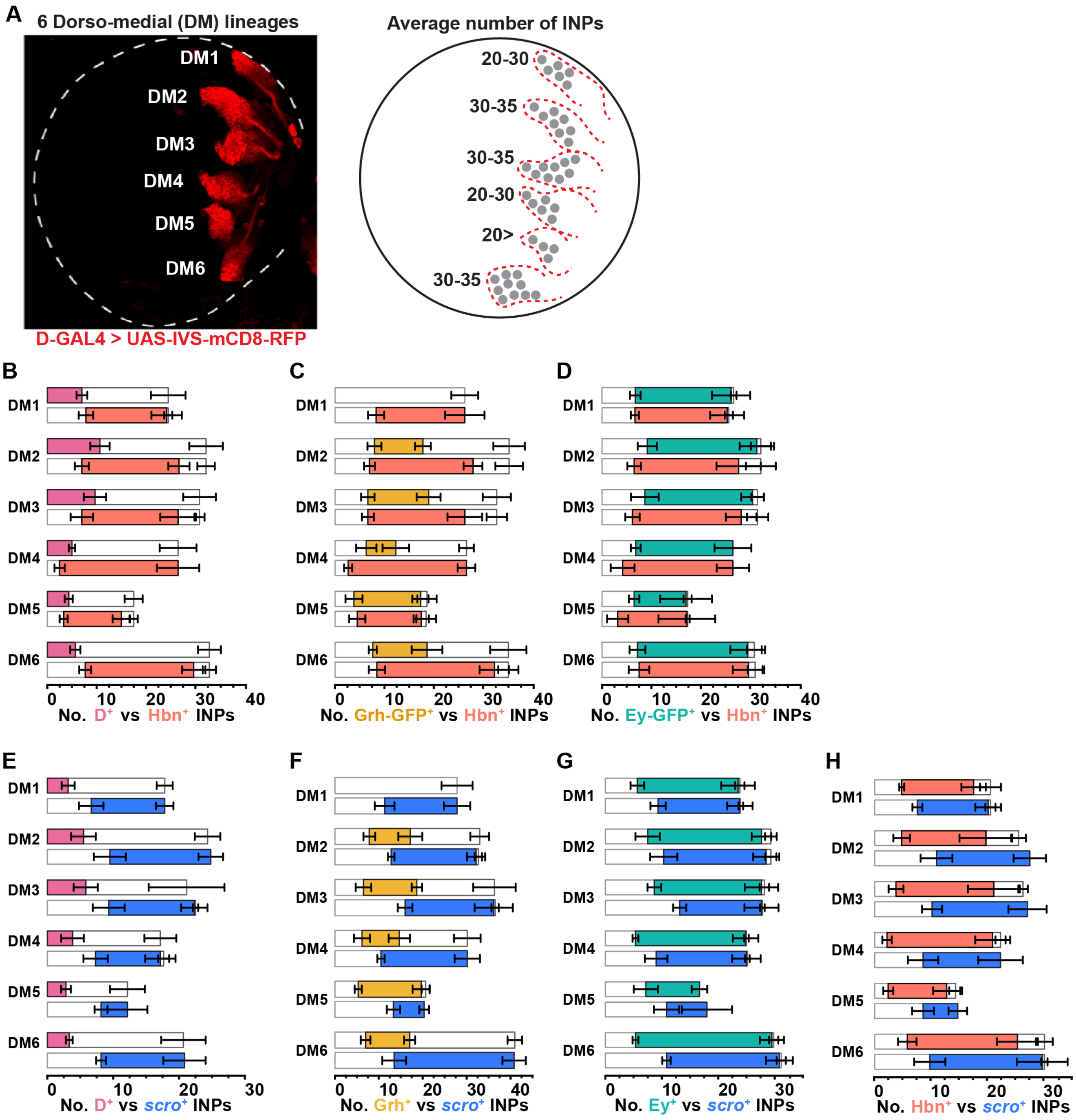
A
6 Dorso-medial (DM) lineages
DM1
DM2
DM3
DM4
DM5
DM6
D-GAL4 > UAS-IVS-mCD8-RFP
Average number of INPs
20-30
30-35
30-35
20-30
20>
30-35
B
C
D
No. D+ vs Hbn+ INPs
No. Grh-GFP+ vs Hbn+ INPs
No. Ey-GFP+ vs Hbn+ INPs
E
F
G
H
No. D+ vs scro+ INPs
No. Grh+ vs scro+ INPs
No. Ey+ vs scro+ INPs
No. Hbn+ vs scro+ INPs

**Figure S3: Overlap of Homeobrain and *scarecrow* with D, Grh and Ey in INPs in DM1-6 lineages.**

(**A**) Schematic showing the 6 dorso-medial (DM) lineages and average number of INPs per lineage.

Quantifications showing:

(**B**) D/Hbn overlap.

(**C**) Grh/Hbn overlap.

(**D**) Ey/Hbn overlap.

(**E**) D/*scro* overlap.

(**F**) Grh/*scro* overlap.

(**G**) Ey/*scro* overlap.

(**H**) Hbn/*scro* overlap.

Single section confocal images. White dotted lines indicate *D*-GAL4>*mCD8-RFP*. Brains were dissected at wandering third instar stage. *n* = 6 brain lobes for all conditions.

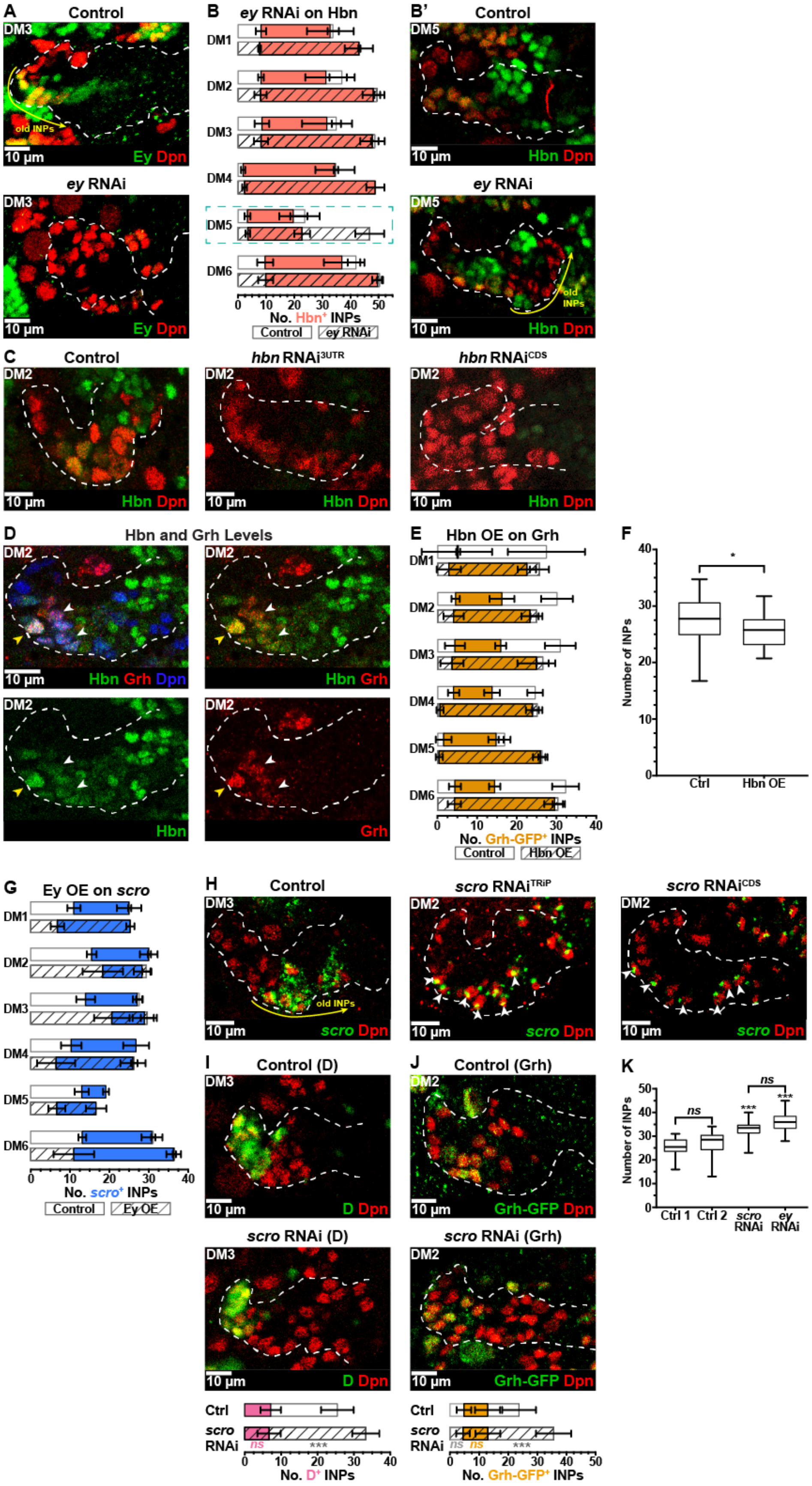

A Control
ey RNAi
Ey Dpn
old INPs
10 µm
B ey RNAi on Hbn
DM1 DM2 DM3 DM4 DM5 DM6
No. Hbn+ INPs
Control
ey RNAi
B' Control
ey RNAi
Hbn Dpn
C Control
hbn RNAi3UTR
hbn RNAiCDS
D Hbn and Grh Levels
Hbn Grh Dpn
Hbn Grh
Hbn
Grh
E Hbn OE on Grh
No. Grh-GFP+ INPs
Hbn OE
F
Number of INPs
Ctrl
Hbn OE
G Ey OE on scro
No. scro+ INPs
Ey OE
H Control
scro RNAiTRiP
scro RNAiCDS
scro Dpn
I Control (D)
scro RNAi (D)
D Dpn
J Control (Grh)
scro RNAi (Grh)
Grh-GFP Dpn
K
Ctrl 1
Ctrl 2
scro RNAi
ey RNAi
ns
***
No. D+ INPs
Ctrl
scro RNAi

**Figure S4: Regulatory Relationships Supplementary.**

(**A**) Control ($w^{1118}$) DM lineages expressed Ey (green) in old INPs (Dpn$^{+}$ (red), yellow arrow). Driving *ey* RNAi resulted in the loss of Ey in INPs. RNAi expression was driven by *D*-GAL4>*mCD8-RFP* in combination with UAS-*Dcr2*. Images are projections over 8 µm in z.

(**B-B'**) Quantification of the effect of *ey* RNAi on Hbn expression in INPs. *ey* RNAi extends the Hbn temporal window in all lineages except DM5. Control line is $w^{1118}$.

(**C**) Driving *hbn* RNAi$^{3UTR}$ or *hbn* RNAi$^{CDS}$ in INPs resulted in the loss of Hbn (green) indicating effective knockdown.

(**D**) The strength of Hbn and Grh expression levels correlate with each other. Yellow arrows indicate strong Hbn and Grh expression, white arrows indicate weaker expressions in INPs.

(**E**) Quantification of Hbn overexpression on Grh expression in INPs in all DM lineages. Control line is $w^{1118}$.

**(F)** Overexpression of Hbn leads to a decrease in INP number. Mann-Whitney Test p=0.041, *.

(**G**) Ey misexpression precociously activates *scro* in INPs in all lineages except DM2 and DM3.

(**H**) Expressing a control RNAi (*mCherry* RNAi) did not affect the expression of *scro* (green) in INPs (Dpn$^{+}$, highlighted by yellow arrow). Driving *scro* RNAi$^{TRiP}$ or *scro* RNAi$^{CDS}$ in INPs resulted in the loss of *scro* mRNA from the cytoplasm of old INPs (arrowheads) indicating effective knockdown. Note that *scro* mRNA can still be detected in the nucleus even when driving *scro* RNAi.

(**I**) *scro* RNAi does not affect D expression in INPs. Control line is *mCherry* RNAi. Mann-Whitney Test p=0.36, ns; p<0.001, ***.

(**J**) *scro* RNAi does not affect Grh expression in INPs. Control line is *mCherry* RNAi. Mann-Whitney Test p=0.26, ns; p=0.41, ns; p<0.001, ***.

(**K**) Knockdown of *scro* or *ey* leads to an increase in the number of INPs. Quantification of the number of INPs in control DM lineages compared to *scro* RNAi or *ey* RNAi lineages. Control 1 is *mCherry* RNAi, Control 2 is $w^{1118}$ and *scro* RNAi is *scro* RNAi$^{TRiP}$. Kruskal-Wallis Test with Dunn's Test. Ctrl1 vs *scro* RNAi: p<0.001, ***; Ctrl1 vs *ey*

RNAi: $p<0.001$, ***; Ctrl2 vs *scro* RNAi: $p<0.001$, ***; Ctrl2 vs *ey* RNAi: $p<0.001$, ***; *scro* RNAi vs ey RNAi: $p>0.99$, ns

Single section confocal images. White dotted lines indicate *D*-GAL4>*mCD8-RFP* expression. Brains were dissected at wandering third instar stage. *n* = 6 brain lobes for all experiments.

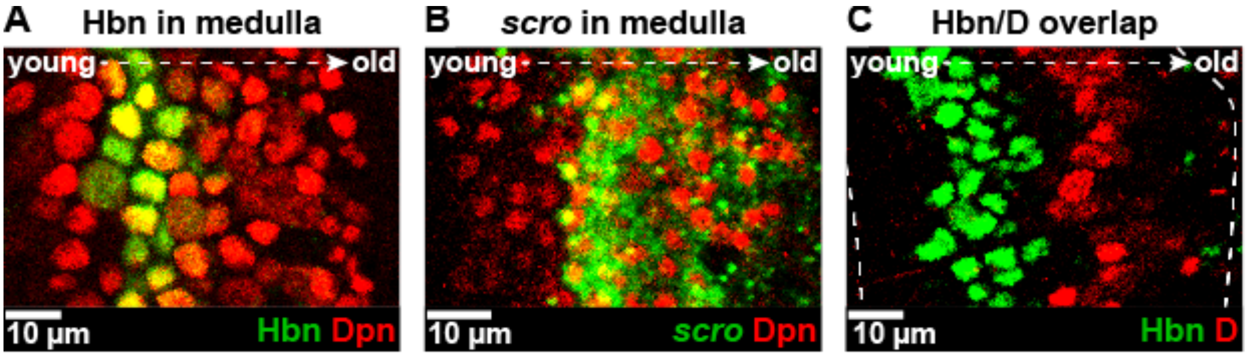


**Figure S5: Hbn and *scro* as optic lobe temporal factors.**

(**A**) Hbn (green) is expressed in a subset of the young OL medulla NSCs (Dpn$^+$ (red)).

(**B**) *scro* (green) is expressed in the old OL medulla NSCs (Dpn$^+$).

(**C**) Hbn (green) and D (red) do not overlap in the OL medulla and the Hbn temporal window precedes the D window.
